# CryoEM reveals how the small molecule EGCG binds to Alzheimer’s brain-derived tau fibrils and initiates fibril disaggregation

**DOI:** 10.1101/2020.05.29.124537

**Authors:** PM Seidler, DR Boyer, MR Sawaya, P Ge, WS Shin, MA DeTure, DW Dickson, L Jiang, DS Eisenberg

**Affiliations:** Departments of Chemistry and Biochemistry and Biological Chemistry, UCLA-DOE Institute, Molecular Biology Institute, and Howard Hughes Medical Institute, UCLA, Los Angeles CA; Department of Neurology, David Geffen School of Medicine, UCLA, Los Angeles CA 90095; California NanoSystem Institute, UCLA, Los Angeles, CA, USA; Department of Neuroscience, Mayo Clinic, Jacksonville, Florida

## Abstract

EGCG, the most abundant favanol in green tea, is one of the few natural compounds known to inhibit amyloid fibril formation of proteins associated with neurodegeneration, and to disaggregate amyloid fibrils. Little is known of the mechanism of molecular action of EGCG, or how it or other small molecules interact with amyloid fibrils. Here we present a 3.9 Å resolution cryoEM structure that reveals the site of EGCG binding to Alzheimer’s disease (AD) brain-derived tau fibrils. The structure suggests that EGCG disaggregates fibrils of AD-tau by wedging into a cleft that is at the interface of two protofilaments of the paired helical filament, and by causing charge repulsions between tau layers of the fibril. In support of this, we observe separation of the protofilaments that EGCG wedges between, and accompanying displacement of the adjacent β-helix domain. By resolving the site of EGCG binding, our structure defines a pharmacophore-like cleft in the AD-tau fibril that will be of use for the discovery of surrogate compounds with more desirable drug-like properties.

## Introduction

Single-particle cryoEM has delivered spectacular structures of protein amyloid fibrils^1–8^, including several of Alzheimer’s associated tau purified from human brain tissues (AD-tau)^1,9,10^. These structures offer the opportunity to learn atomic level details of how known inhibitors interact with amyloid fibrils. Epigallocatechin gallate (EGCG), a small molecule natural product in green tea, inhibits aggregation of numerous amyloid proteins and, despite their tremendous stability, disaggregates fibrils of some amyloids into smaller units.11-15 Evidence suggesting that EGCG disaggregates fibrils of tau includes studies in primary rat neurons showing that EGCG enhances clearance of phosphorylated tau,^16^ although the reported effects of EGCG are somewhat pleiotropic and range from the inhibition aggregation to altered post-translational modification.^17,18^. Here, we sought to illuminate the effects direct binding of EGCG to AD-tau by single-particle cryoEM structure determination. The challenge is to obtain a picture of the kinetic intermediate of pathogenic tau fibrils bound to EGCG, yet prior to the subsequent disaggregated state.

Aggregated tau forms fibrillar structures that propagate and cause neurodegeneration by prion-like seeding.^19,20^ Single-particle cryoEM structures of brain-purified tau from donors with AD show that pathological tau fibrils, like other pathogenic amyloids, are extraordinarily stable owing to steric zipper interactions, and an extensive network of polarized hydrogen bonds (H-bonds) that forms along the fibril axis.^9,10,21,22^ The binding enthalpies of resulting amyloid fibril structures rivals that of crystalline ice.^23^ Owing to these features, pathogenic amyloid fibrils can withstand heating to boiling temperatures and resist solubilization in detergents such as sarkosyl, explaining the ability of parent fibrils to seed new fibrils and to evade degradation by cellular machinery; and at the same time, presenting the question of how amyloid fibrils can be disaggregated by a small molecule.

Amyloid fibrils are disaggregated biologically by the aid of specific chaperones proteins,^24–27^ and/or the autophagosome^16^. Amyloid disaggregation is an attractive potential therapeutic mechanism of action.^13,28^ Whereas chaperones function as high molecular weight homo- and heteromeric assemblies that use chemical energy to disaggregate yeast prions and certain amyloid fibrils^,2^5,26 the mechanistic details of amyloid fibril disaggregation by small molecules is more mysterious. Remarkably, the small molecule natural product EGCG is reported to inhibit at least 14 different amyloids, in part by disaggregating amyloid fibrils, although also by blocking aggregation of protein monomers.^11–13,15,29^

Despite potent in vitro anti-amyloid activity, EGCG is not a proven clinical therapeutic. The lack of clinical efficacy of EGCG stems from poor drug-like properties, which includes limited bioavailability and lack of specificity.^28^ In addition, EGCG is known to interact with numerous biomolecules, often with differing ligand poses^30–34^, and has been described as a pan-assay interfering compound.^35^ Reflecting its ability to bind promiscuously to proteins with differing structures, EGCG is reported to modulate the activity of dozens of different disease-related pathways and proteins^36^ and broadly inhibits amyloid proteins.^11^ Despite EGCG’s limitations as a clinical agent, we sought to understand how EGCG interacts with, and inhibits the AD-tau fibril polymorph. The EGCG binding site we discovered on the AD-tau fibril is quite unlike EGCG binding sites in globular proteins, and constitutes an AD-tau amyloid pharmacophore that can possibly be exploited to create small-molecule inhibitors with drug-like properties.

### EGCG disaggregates tau by stoichiometric binding to fibrils

To confirm that EGCG binds to amyloid fibrils of tau, and that inhibition of tau by EGCG is not the result of indirect effects,^16–18^ we confirmed binding of EGCG to recombinant fibrils of tau by ITC (Fig. 1a). We found that EGCG binds to recombinant tau-K18+ fibrils with an affinity of 1.6 μM and near equimolar stoichiometry, with an N of 0.86.

**Figure 1.**
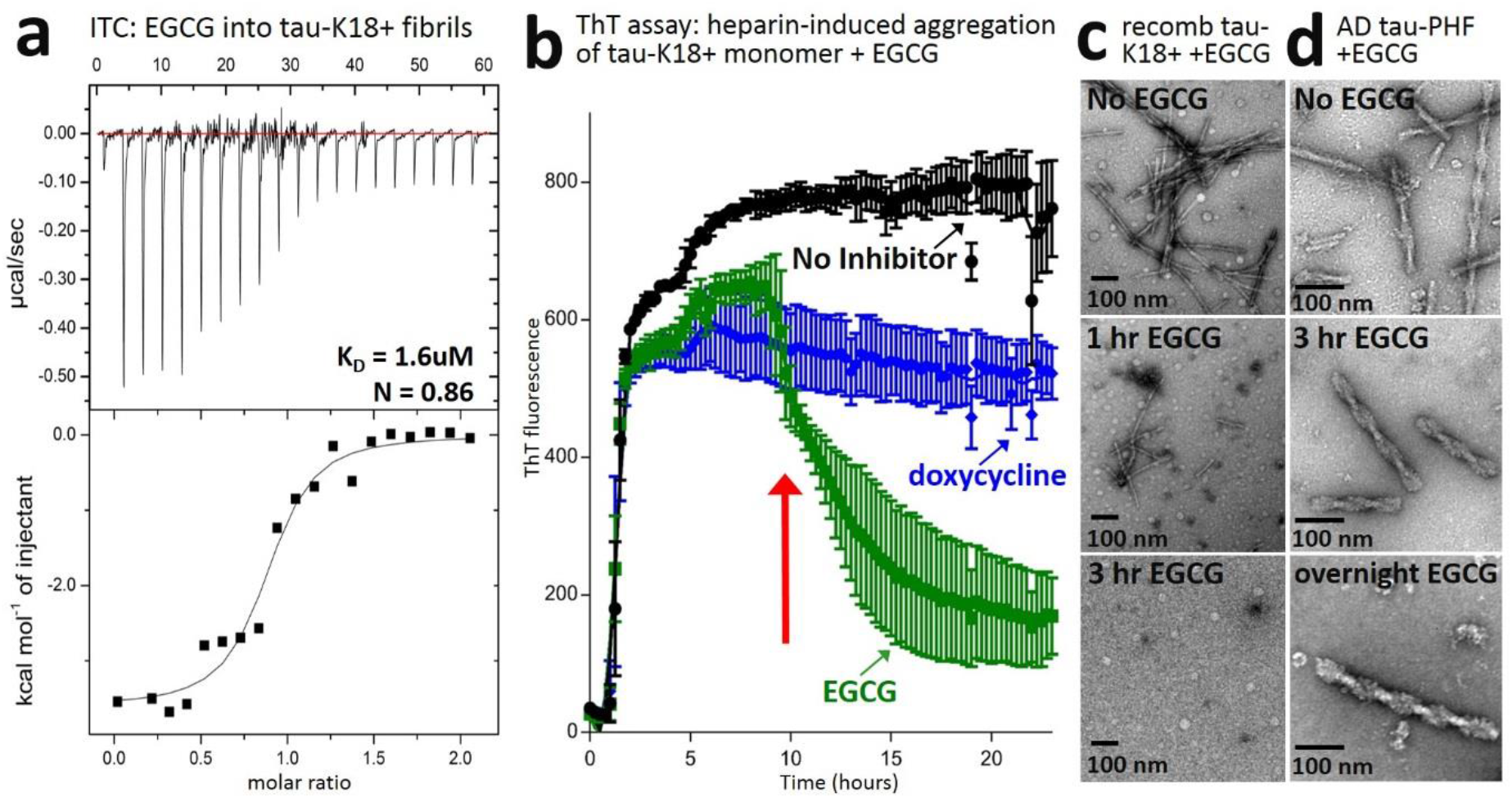
EGCG binds to and disaggregates tau fibrils. (a) Isothermal titration calorimetry (ITC) of 750 μM EGCG into 75 μM recombinant tau-K18+ fibrils, a construct that contains all of the residues that are observed in the AD-tau fibril core38. The binding isotherm was fit using a standard one site binding model. (b) ThT assay showing the effect EGCG or negative control, doxycycline, have on the aggregation of 20 μM tau-K18+ monomer. Note the sudden drop in ThT signal at 10 hrs (marked by a red arrow) for the sample containing EGCG. Results are representative of data from three independent experiments. (c) Negative stain electron micrographs of 75 μM recombinant tau-K18+ fibrils prior to incubation with EGCG (top), or following 1 or 3 hour incubation with a two-fold molar excess of EGCG at room temperature, as indicated. (d) As in c, except EGCG micrographs of tau are of fibrils purified from AD brain tissue and EGCG treatments were carried out at a concentration of 100 μM for a 3 hour or overnight duration at 37 °C, as indicated.

To examine the effect of EGCG on fibril formation of recombinant tau, we carried out the thioflavin T assay shown in Fig. 1b. We observed no shift in the lag time of tau aggregation, suggesting that EGCG has no inhibitory effect on the tau-K18+ monomer. However, fibrils formed in the presence of EGCG were metastable insomuch as the ThT signal abruptly fell after reaching a plateau at 10 hr (Fig. 1b), suggesting that EGCG binding either causes tau fibrils to disaggregate or displaces the ThT dye from the fibrils. Consistent with the hypothesis that EGCG disaggregates fibrils of tau, negative staining electron microscopy shows a time-dependent disappearance of recombinant tau fibrils following the addition of EGCG (Fig. 1c) with a large fraction of fibrils disappearing within 1 hr of the addition of EGCG, and virtually all fibrils having disappeared by 3 hrs. An alternative explanation that EGCG prevents fibrils from adhering to the EM grid is unlikely because the disappearance of fibrils becomes gradually pronounced over time in spite of all other variables remaining unchanged, and also because others report that EGCG disaggregates fibrils of other amyloids, including α-synuclein, amyloid β, and apolipoprotein A-I.^12,15^

The effect of EGCG treatment on AD-tau could differ from recombinant tau fibrils since recombinant fibrils of tau form different polymorphs. To test whether EGCG disaggregates brain-purified AD-tau fibrils, we examined negatively stained EM grids of purified AD-tau at various times following incubation with EGCG. AD-tau fibrils were more robust than recombinant tau fibrils and disaggregation was accelerated by incubation at 37 °C. As shown in Fig. 1d before incubation with EGCG, AD-tau fibrils appear as relatively narrow twisted filaments with a high abundance on the EM grid. A 3 hr incubation with EGCG produced fibrils with radically different characteristics compared to non-inhibitor treated fibrils (compare Fig. 1d top and middle), which were broadened and more braided in appearance. Following overnight incubation with EGCG, most of AD-tau fibrils were lost; although some remain, unlike recombinant tau fibrils, which virtually vanish after a 3 hr incubation with EGCG. The AD-tau fibrils that remained after overnight incubation with EGCG appeared broadened and overtly shabby (Fig. 1d, bottom). We conclude that the EGCG has similar disaggregating effects on recombinant and AD-tau fibrils, although it is not likely that recombinant and AD-tau share a structurally conserved EGCG binding cleft given the differences that are reported for recombinant and brain-derived fibrils.^9,10,37^ Given EGCG’s ability to bind polymorphically in protein cavities with diverse shapes and chemical features, it is more likely that EGCG acts on recombinant and AD-tau by interacting with different binding cavities on the two polymorphs.

### The EGCG binding cleft on fibrils of AD-tau

To identify the binding cleft of EGCG on the AD-tau fibril polymorph, we turned to single-particle cryoEM since structures of AD-tau are exceptionally conserved among five different brain donors from which structures were previously determined by labs around the world.^1,9,10^ We purified AD-tau using the same protocol detailed by authors of these reports. To resolve an intermediate conformation of AD-tau bound to EGCG (one that precedes full disaggregation and retains sufficient order to inform about the action of EGCG), we pre-incubated purified AD-tau fibrils with EGCG for 3 hr at 37 °C immediately prior to freezing cryoEM grids. These conditions yielded fibrils with overall similarity to non-EGCG treated fibrils as judged by negative-stain EM (Fig. 1d) but with new features revealed by cryoEM.

We determined the structure of AD-tau fibrils that were pre-incubated with EGCG by helical reconstruction using unbiased 3D classifications of manually picked particles with a cylindrical reference having an outer diameter of 250 pixels (266 Å). The only AD-tau polymorph that emerged from 3D reconstructions was the PHF, suggesting that the other known AD-tau fibril polymorph, straight filaments, were either particularly sensitive to treatment with EGCG, or too few to observe. It is noteworthy that straight filaments are the minor fibril polymorph of AD-tau.^9,10^

As shown in Fig. 2a, the spacing between individual β-strands (having a known distance of 4.8 Å) was apparent in the 3D projection of aligned particles extracted using a 686-pixel box, consistent with our resolution estimate of 4.4 Å (determined using a 0.5 FSC cutoff, see Methods). As added evidence of the quality of our EM map, we observed conserved densities that are seen in the five other published AD-tau PHF structures;^1,9,10^ specifically: (1) islands of density proposed to derive either from ubiquitinatio^n1^ or the N-terminal segment, 7-EFE-9, around residues K317 and K321^10,39^, (2) a large patch of density near Q307, V309 and K311 thought to be ubiquitin1 and/or steric zipper interaction with other tau sequences^38^, and (3) two patches of density in the interior core of the fibril (Fig. 2b and c, labels 1-3 in red text).

**Figure 2.**
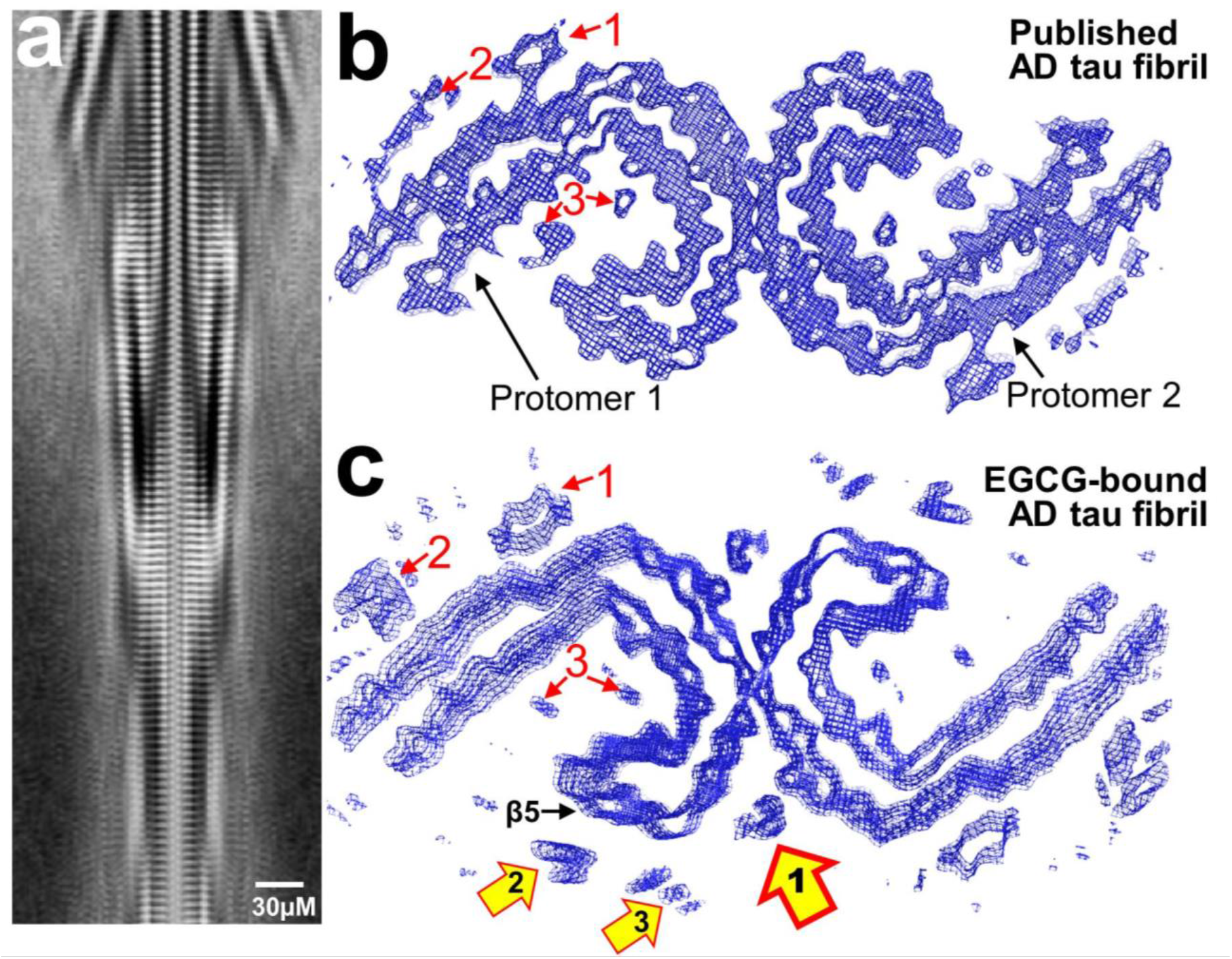
Transient cryoEM structure of AD brain-derived tau fibril bound to inhibitor, EGCG. (a) Cross-section of fibril from 3D classification showing strand separation, with a spacing of 4.8 Å. (b) Published map of AD-brain derived tau PHF (PDB 5O3L), viewed down the fibril axis, for comparison with c. Conserved patches of density are labeled with red text and arrows. See text for additional explanation. (c) Cryo-EM map of AD-brain derived tau PHF complexed with EGCG with an estimated resolution of 4.4 Å. Yellow arrows in c indicate new densities that are attributed to bound EGCG. The location of β-sheet 5, formed by residues Lys343 to Asp348, is labeled β5.

In addition to the conserved patches of density, we observed three new peaks highlighted by yellow-filled arrows in Fig. 2c that we attribute to bound EGCG. Peak 1 lies in a cleft that is formed by the junction of two tau protofilaments, and is found at both symmetry related sites of the PHF. Peaks 2 and 3 lie on the surface of the fibril, and could represent additional sites of EGCG binding sites. Both Peaks 2 and 3 are in proximity to lysines that are part of a β-helix that is, as we discuss in detail below, evidently mobilized by EGCG binding (Fig. 4 and 3b)—Peak 2 is nearest Lys347 and Peak 3 is nearest Lys343. We also consider it an alternate possibility that Peaks 2 and 3 arise from protein densities that reflect a subpopulation of molecules that have undergone a conformational change that was induced by EGCG-binding and the 3 hr pre-incubation prior to freezing EM grids. In the case of the latter hypothesis, it is possible that Peaks 2 and 3 arise from the translation of β-sheet 5, formed by residues Lys343 to Asp348, away from the fibril’s center of mass. This hypothesis is supported by the fact that unlike Peak 1, neither Peaks 2 nor 3 foster a chemical environment that is typical of other ligand binding sites—both are highly solvent-exposed, and owing to their remote nature on the protein backbone, offer few protein contacts for binding. However at the resolution of our map, we cannot determine with confidence the origin of Peaks 2 and 3.

**Figure 3.**
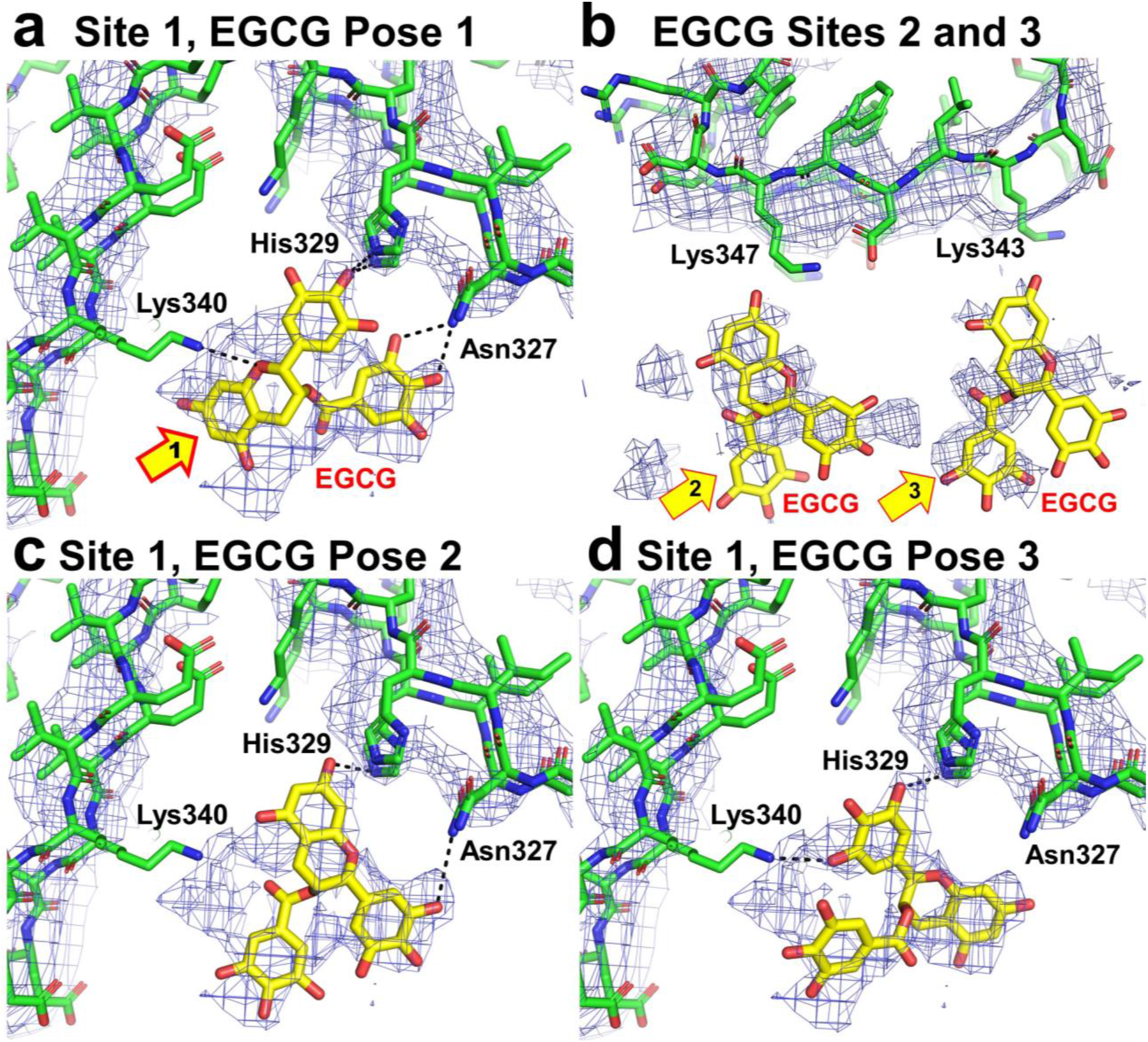
Models of possible binding poses of EGCG in cryoEM densities of AD-tau fibrils. (a) EGCG fit in Peak 1 density, shown in Fig. 2c. (b) EGCG fit in densities Peak 2 and 3 from Fig. 2c. (c and d) Alternative EGCG poses fit in Peak 1 density, as in a.

**Figure 4.**
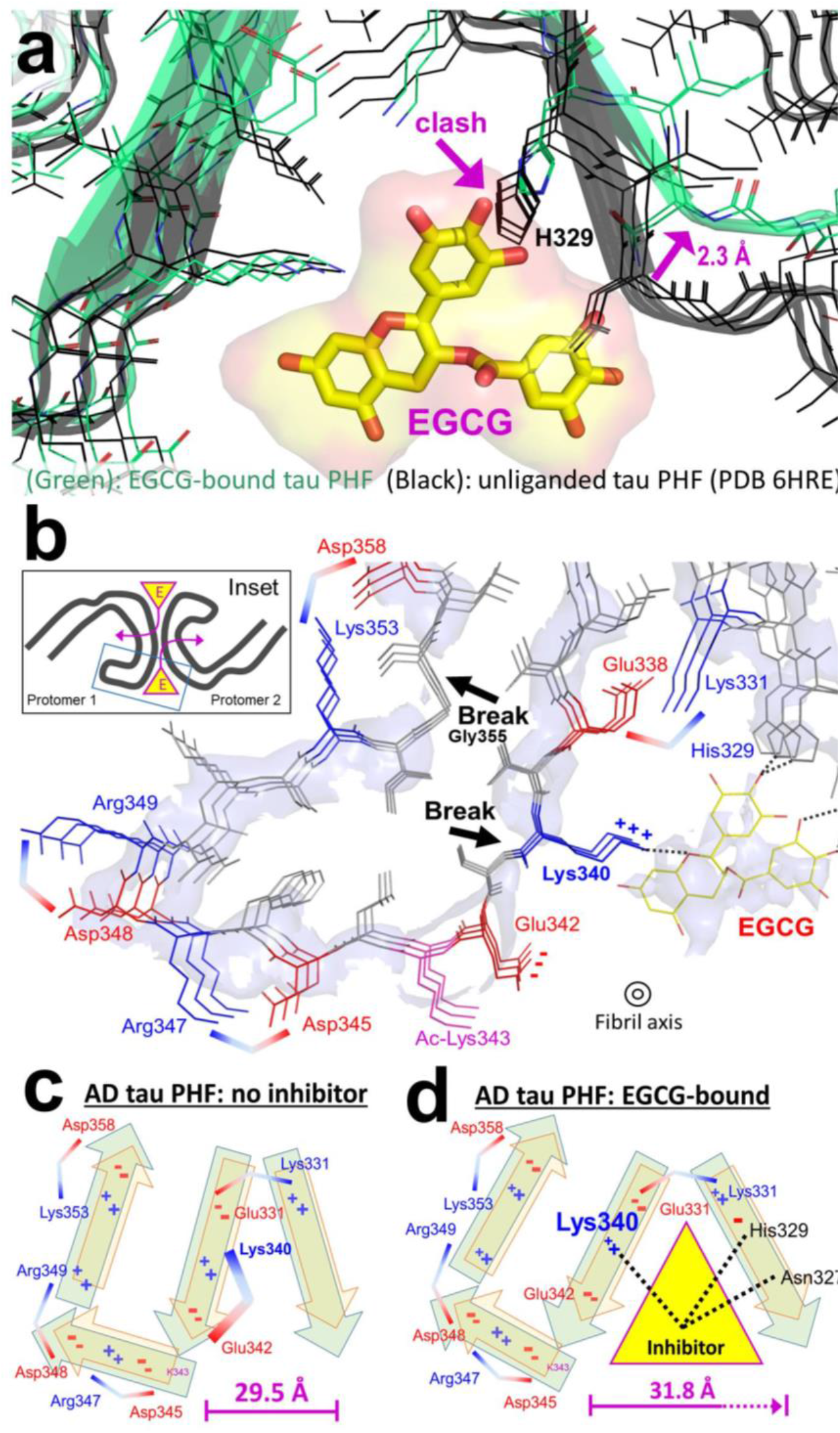
The wedge-charge-destabilization hypothesis for fibril disaggregation by EGCG. (a) Overlay of EGCG-bound (green) and unliganded (black, PDB 5O3L) tau PHFs showing 2.3 Å separation of tau protofilaments in the EGCG-bound form, and imposed clash of EGCG with His329 in the unliganded conformation. EGCG is shown as yellow/red space fill. (b) Alternating negatively (red) and positively (blue) charged sidechains of the β-helix segment of AD-tau shown from the enlargement of the boxed region of the Inset. Stabilization of the fibril structure is provided by pairing of neighboring oppositely charged residues (pairs shown by boomerang-shaped brackets). By forming a H-bond with Lys340 of tau (dotted black line), bound EGCG diminishes the effect of ion pair-mediated charge stabilization of positive Lys340 with its neighboring negative Glu342, thereby increasing inter-layer repulsion with neighboring layers. This repulsion would be expected to weaken inter-layer bonding, favoring fibril disruption. (c) Schematic of the unliganded PHF viewed down the fibril axis, and (d) schematic of the wedge effect of EGCG, forcing the two protofilaments apart, and disrupting ion paired residues at the EGCG binding interface.

To visualize in better detail the contacts of EGCG with the AD-tau fibril, we obtained a higher resolution map by re-extracting the high resolution subset of particles from the 686-pixel box boxes with smaller box sizes. 3D refinement using a 432-pixel box size produced a 3.9 Å map (determined using a 0.5 FSC cutoff, see Methods and Supplementary Fig. 1), and Postprocess analysis performed with Relion suggests the 3.9 Å map is of good quality with no evidence of overfitting. We suspect that our inability to obtain particle alignments with even smaller box sizes for higher resolution map calculations was limited by structural heterogeneity that is induced by the 3 hr pre-incubation of AD-tau with EGCG prior to freezing. Whereas higher resolution structures are now common for single-particle cryoEM studies of well-ordered substances, capturing an image of a fibril in the process of disaggregation limits attainable detail.

We used our 3.9 Å resolution map to model EGCG into the 3 peaks of density that we discovered from our low-resolution map. As shown in Fig. 3, EGCG could be modeled into the density of Peak 1 with the greatest confidence. The best pose of EGCG that was modeled in Site 1 (Fig. 3a) binds between layered protofilaments of tau making a total of 5 potential H-bonds with 3 residues from AD-tau: the hydroxyl of one monocyclic ring of EGCG is within H-bonding distance of two His329 residues (1 from each protomer in the nearest layers above and below the molecule of bound EGCG. In addition, hydroxyls at the *ortho*- and *meta*-positions of the other monocyclic ring of EGCG are within H-bonding distance of Asn327 of the same protofilament. An additional potential H-bond is made between Lys340 from the opposite tau protofilament and the oxygen of the ether in the bicyclic moiety of EGCG.

Our analysis of the various modeled EGCG ligands, shown in Fig. 3, supports our conclusion that EGCG binds to AD-tau predominantly with a single best inhibitor pose that is superior to alternative competing inhibitor binding poses that we considered. Molecules of EGCG that are docked in Sites 2 and 3, corresponding to Peaks 2 and 3 from Fig. 2c, form fewer protein contacts owing to the remote site of binding, and the considerable distance of Peaks 2 and 3 from the protein backbone (Fig. 3b). For these reasons, the intensity and quality of Peaks 2 and 3 are of notably poorer quality than Peak 1, and as a result, we cannot conclude with certainty whether or not Sites 2 and 3 are bonafide sites of EGCG binding.

The EGCG binding pose shown in Fig. 3a is likely the primary mode of EGCG interaction with AD-tau although at 3.9 Å, the resolution of our best cryoEM map, it is impossible to determine the exact hydrogen bonds that are made by EGCG with AD-tau based on fitting alone. To guide our analysis, we modeled alternative EGCG binding poses in Peak 1 density of the Site 1 EGCG binding cleft (Fig. 3c and d). Both alternative EGCG binding poses that we modeled, Poses 2 and 3, make plausible H-bond contacts with AD-tau; Pose 2 with Asn327 and His329, and Pose 3 with His329 and Lys340 of the opposite protomer. All of the modeled EGCG poses fit portions of the Peak 1 density, although none make as many stabilizing contacts as Pose 1. We conclude that alternate poses of EGCG, potentially at partial occupancy, are compatible with the Site 1 cleft.

### Effects of EGCG on the structure of AD-tau

Examination of the 3.9 Å resolution density shows changes in AD-tau induced by the binding of EGCG. EGCG appears to wedge into the Site 1 EGCG binding cleft, pushing the AD-tau dimer apart by ~2.3 Å (Fig. 4a). This minor movement of the protein backbone is of major significance because it relieves a steric clash between EGCG and His329. Accompanying the backbone movement around the Site 1 EGCG binding cleft are breaks in the backbone density at residues Lys340 and Gly355 of the neighboring tau protomer. We attribute these breaks to the partial displacement of a hinge region that is mobilized by the wedging effect of EGCG, rather than poor map quality. That is, we interpret these breaks to be biochemically meaningful disruptions that offer a glimpse of the strain in the fibril that is induced by EGCG just prior to disaggregation. This hypothesis is supported by the proximity of the broken densities to the EGCG binding cleft, and these being the only breaks in backbone density in the entire structure.

## Discussion

The EGCG inhibitor binding cleft on the tau PHF is quite unlike binding sites of small molecules on globular and membrane proteins^30–34^. Firstly it is not a single site, but rather is a contiguous column that is created by thousands of stacked inhibitor binding clefts. Each bound EGCG molecule is close enough to EGCG molecules in the layers above and below to form self-stabilizing π-stacking interactions by all three rings of the stacked inhibitors. Secondly, each of the many EGCG binding clefts bristles with charged and polar sidechains, ready to form hydrogen bonds with some of the 11 hydrogen-bond acceptors, and 8 hydrogen-bond donors on EGCG. Fig. 4b shows the charged sidechains on the protofilament to the left of EGCG; these include Glu338, Lys340, and continuing along the β-helix Glu342, Lys343, Asp345, and Arg347. Notice that the charges of these sidechains alternate negative and positive. In fact, the pattern of alternating acids and bases makes the PHF structure possible: in order for like charges to stack roughly 4.8 Å apart along the fibril axis, neighboring residues on the same level can form ion pairs to counteract charge repulsion between identical residues in adjacent layers. This arrangement of alternating acids and bases, which we refer to as intra-layer charge stabilization, was observed in the previous high-resolution structure of the CTE tau polymorph.4 Perhaps also helping to balance charges are bound water molecules and/or post-translational modifications such as the acetylation of Lys3431. Potential hydrogen bonding residues on the other protofilament include Asn327, His329, and Lys331.

Our Wedge-Charge-Destabilization hypothesis is summarized by Figures 5b-d. By rotation about its single bonds, EGCG adopts a roughly planar wedge to fit into the wedge-shaped clefts at the junctures of the two protofilaments of the PHF of tau. EGCG, by being in position to form hydrogen bonds with Lys340, His329, and Asn327, diminishes intra-layer charge stabilization, thereby increasing inter-layer charge repulsion of stacked charged residues. For example, by H-bonding with Lys340, EGCG disrupts ion pairing with neighboring acidic residues, and increases inter-layer repulsion by creating stacks of unpaired lysines. Lys340 is particularly susceptible to the effect of charge destabilization since it is solvent excluded by the protofilament interface, and is increasingly excluded from solvent by the column of bound EGCG molecules. The disruption of ion pairing in the solvent excluded environment of amyloid fibrils is found to be detrimental to aggregation, whereas outward pointing residues with unpaired charge appear to be somewhat more tolerated.^40^

EGCG binding at the protofilament junction also destabilizes the tau PHF by forcing apart the two protofilaments that form the dimer interface (Fig. 4b and c). The wedge effect of EGCG binding is apparent both from the translation of the protofilament backbone adjacent to the EGCG binding cleft, shown in Fig. 4a, and from breaks in the density at Lys340 and the hinge point of Gly355 (Fig. 4b). These are the only breaks in density that are observed in the structure of the EGCG complex, and suggest that EGCG exerts its effect primarily by perturbing the β-helix domain. In short, we attribute the dissolution of fibrils by EGCG to the combined wedge effect of forcing the protofilaments apart, and the inhibition of charge stabilization, which weakens the binding of layers to one another.

In summary, the structure of EGCG bound to AD-tau fibrils illustrates how an exceptionally rich H-bonding molecule that nestles between protofilaments of amyloid fibrils can weaken interactions that otherwise stabilize fibrils. At the same time, our structure agrees well with the consensus that there is no significant variability in AD-tau fibril structures between individuals with AD.^9^ In keeping, we observed all of the same conserved features in the EGCG-bound fibril of AD-tau, including a protomer fold that is essentially identical to all other AD-tau fibril structures determined, with the exception of a few sterically explainable perturbations that can be attributed to bound EGCG. In addition, our structure defines what might be termed a pathogenic amyloid pharmacophore. Previous suggestions of how small molecules interact with and inhibit fibrils have been mainly limited to computational docking. The structure of the pharmacophore of the AD brain-derived tau PHF thus opens the door to informed screening and the design of small molecules with drug-like properties.

## Experimental Methods

### Preparation of crude and purified brain-derived tau seeds

Purification of tau PHFs and SFs from AD brain tissue were carried out exactly as described previously.^10^ Prior to CryoEM grid preparation, AD brain-purified tau fibrils were pre-incubated with 0.5 mM EGCG in 20 mM Tris–HCl pH 7.4, 100 mM NaCl at 37 °C for 3 hrs.

### CryoEsM data collection and 3D reconstruction

AD brain purified tau fibrils were applied to negatively glow-discharged Quantifoil 1.2/1.3 electron microscope grids (2.6 μl for 1 min), and subsequently plunge-frozen in liquid ethane on a Vitrobot Mark IV (FEI). Data were collected on a Titan Krios (FEI) microscope equipped with a Gatan Quantum LS/K2 Summit direct electron detection camera (operated with 300 kV acceleration voltage and slit width of 20 eV). Counting mode movies were collected on a Gatan K2 Summit direct electron detector with a nominal physical pixel size of 1.07 Å per pixel with a dose per frame 1.26 e-/Å2. A total of 30 frames with a frame rate of 5 Hz were taken for each movie, resulting in a final dose 38 e-/Å2 per image. Automated data collection was driven by the Leginon automation software package.^41^

Data were processed in a similar manner as the SegA-sym and SegA-asym structures from Cao, et al.^42^ Briefly, CTF estimation was performed using CTFFIND 4.1.8^43^, and Unblur^44^ was used to correct beam-induced motion. Particle picking was performed manually using EMAN2 e2helixboxer.py^45^, and particles were extracted and classified in RELION^46^ using the 90% overlap scheme. Particles were extracted using a 686-pixel box size for 2D classifications and a first series of 3D classifications. Typically, we performed Class3D jobs with K = 3 and manual control of the tau_fudge factor and healpix to reach a resolution of ~5-6 Å to select for particles that contributed to the highest resolution class. After determining the set of particles that contributed to the highest resolution 686-pixel box size reconstruction, we re-extracted all fibrils from which these particles originated with the same 90% overlap scheme as above. Two subsequent rounds of 3D classification were carried out for the 432-pixel box particles. For the final high-resolution refinement, Refine3D was unable to regulate the working resolution with high enough resolution factors to push past 4.8 Å perhaps owing to low particle numbers, or due to the fact that the particles bound to EGCG are in the process of disassembly and are therefore less well ordered. Therefore, we employed a ‘suboptimal’ manual refinement with a single reference map and all particles, as described previously.^8,47^ To assess the resolution of the final reconstruction, we split the particles contributing to the final map *a posteriori* into even and odd micrographs, and we reconstructed two-half-maps using the fully refined alignment parameters. We then performed the map–map Fourier shell correlation (FSC) using these two half-maps with a generous, soft-edged solvent mask and high-resolution phase randomization in RELION PostProcess to test for over-fitting.^48^ The resulting map has a resolution estimate of 3.9 Å with a 0.5 FSC cutoff with no indication of over-fitting.

### Atomic model building

The highest resolution structure of an AD-brain derived tau PHF (PDB 6HRE) was used as a starting model for atomic model building. Coordinates were docked in the 3.9 Å density map as a rigid body, and minor fitting was performed in COOT^49^. EGCG was modeled into the density by first rigid body docking from molecule KDH 911 of PDB 4AWM and subsequent real space refinement. Fibril layers were modeled to maintain local contacts between chains in the fibril during structure refinement. We performed automated structure refinement using phenix.real_space_refine^50^.

### Negative stain grid preparation

Negatively stained EM Grids were prepared by depositing 6 μl of sample on formvar/carbon coated copper grids (400 mesh) for 3 min with. Sample was rapidly wicked using filter paper without drying the grid, and stained with 1% uranyl acetate for 2 min.

### Isothermal Titration Calorimetry

Fibrils of tau-K18+ were prepared at a concentration of 50 μM in 1X PBS (pH 7.4) with 0.225 mg/ml heparin (Sigma cat. H3393) and 1 mM dithiothreitol with shaking at 700 rpm at 37 °C with the addition of a plastic bead to enhance agitation. A solution volume of 0.5 ml was necessary for each ITC experiment. After 18 hrs, the cloudy solution was pelleted by ultracentrifugation at 60,000 rpm for 30 min and the pellet was resuspended in 0.3 ml of 1X PBS. Fibril formation under these conditions yielded no detectable tau-K18+ in the supernatant of the ultra-spin by SDS-PAGE, so the concentration of tau fibril solution prepared by this method is estimated to be 75 μM. EGCG was dissolved in 1X PBS at a concentration of 750 μM. ITC was carried out by titration of EGCG into tau fibril solution with a MicroCal iTC200 at 23 °C. Binding isotherms were fit conservatively using a standard one site binding model.

### ThT aggregation kinetics assay

Concentrated tau-K18+ was diluted into PBS buffer (pH 7.4) to a final concentration of 20 μM with an equimolar ratio of indicated small molecule inhibitor. Proteins were shaken in solutions containing 40 μM ThT, 0.225 mg/ml heparin and 1 mM dithiothreitol in a 96-well plate with a plastic bead to enhance agitation. ThT fluorescence was measured with excitation and emission wavelengths of 440 and 480 nm, and averaged curves were generated from triplicate measurements as indicated in the figure legend. Error bars show the standard deviation of replicates measurements.

## Acknowledgements

We thank the donors and their families; without whom this work would not have been possible. We acknowledge 1R01 AG029430 (DSE), AG061847 (DSE) and RF1 AG054022 (DSE) from the National Institute on Aging, 1F32 NS095661 from the National Institute of Neurological Disorders and Stroke (PMS), A2016588F from the BrightFocus Foundation (PMS) and HHMI for support.

## Data Availability

Structure coordinates and map files are deposited into the Worldwide Protein Data Bank (wwPDB) and the Electron Microscopy Data Bank (EMDB) with the following accession codes: PDB ID 6W9B and EMD-21581.

## Supplementary Figures

**Supplementary Figure 1.**
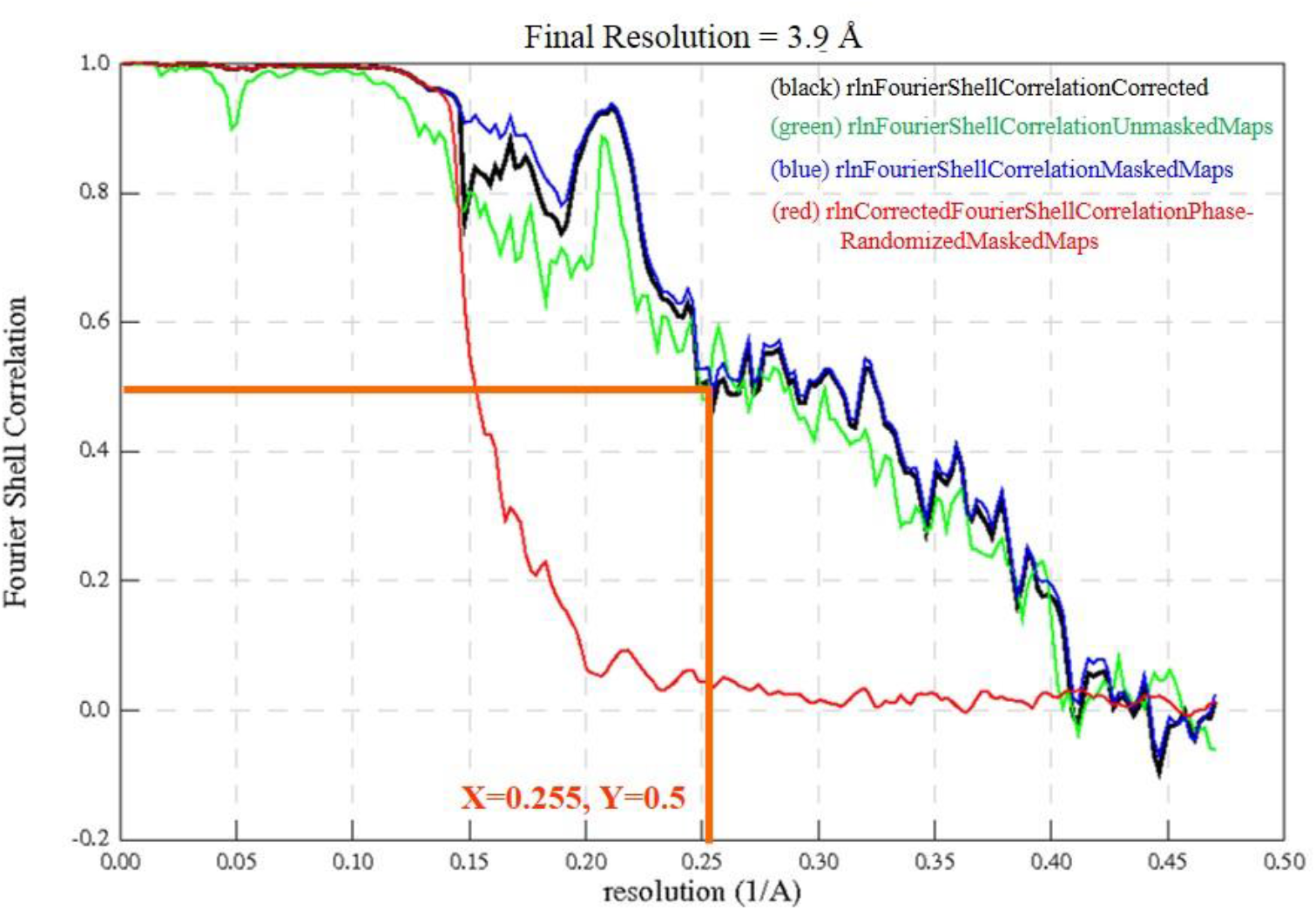
Fourier Shell Correlation (FSC) plot used to determine resolution of 432-pixel data set using a 0.5 FSC cutoff, as indicated with orange lines. FSC analysis was performed using postprocess in Relion^46^, and confirms lack of overfitting owing to strong agreement between the unmasked (green), masked (blue) and corrected black) curves. In addition, the curve plotted for the phase randomized masked map (red) shows sharply approaches zero indicating lack of correlation of noise between two half maps, also suggesting maps lack indication of overfitting.

**Table 1.**
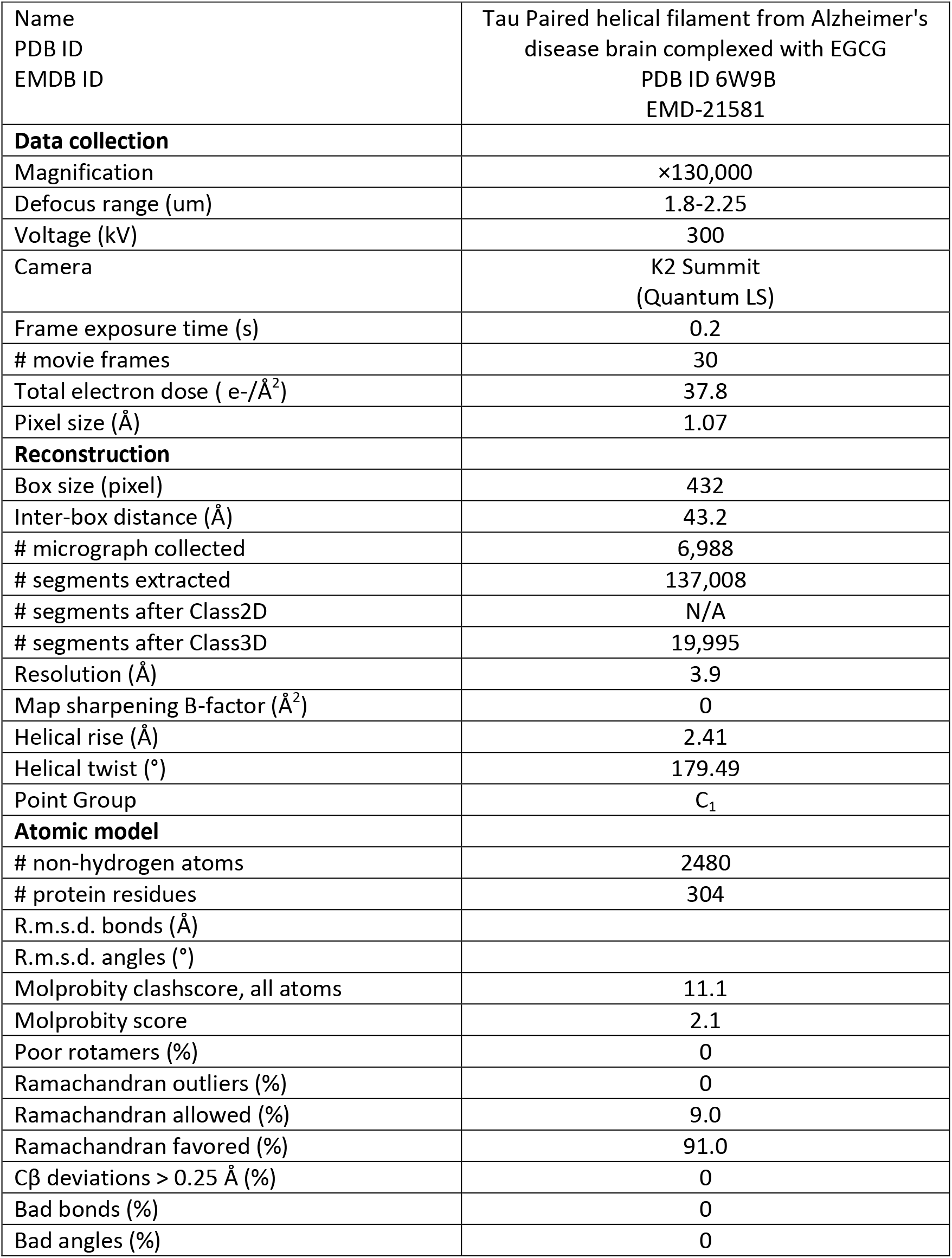
Cryo-EM data collection, refinement, and validation statistics.

