## Supplement for "CryoEM reveals how the small molecule EGCG binds to Alzheimer’s brain-derived tau fibrils and initiates fibril disaggregation"

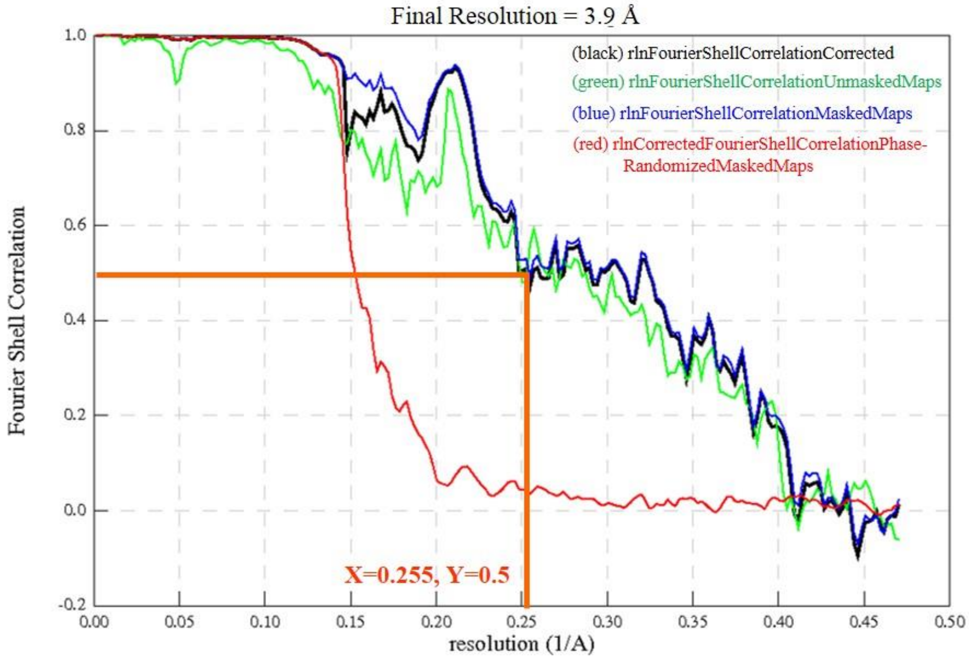


**Supplementary Figure 1** Fourier Shell Correlation (FSC) plot used to determine resolution of 432-pixel data set using a 0.5 FSC cutoff, as indicated with orange lines. FSC analysis was performed using postprocess in Relion[46], and confirms lack of overfitting owing to strong agreement between the unmasked (green), masked (blue) and corrected black) curves. In addition, the curve plotted for the phase randomized masked map (red) shows sharply approaches zero indicating lack of correlation of noise between two half maps, also suggesting maps lack indication of overfitting.

Table 1 Cryo-EM data collection, refinement, and validation statistics.

| | |
|---|---|
| Name<br>PDB ID<br>EMDB ID | Tau Paired helical filament from Alzheimer's disease brain complexed with EGCG<br>PDB ID 6W9B<br>EMD-21581 |
| **Data collection** | |
| Magnification | ×130,000 |
| Defocus range (um) | 1.8-2.25 |
| Voltage (kV) | 300 |
| Camera | K2 Summit<br>(Quantum LS) |
| Frame exposure time (s) | 0.2 |
| # movie frames | 30 |
| Total electron dose ( e-/Å$^2$) | 37.8 |
| Pixel size (Å) | 1.07 |
| **Reconstruction** | |
| Box size (pixel) | 432 |
| Inter-box distance (Å) | 43.2 |
| # micrograph collected | 6,988 |
| # segments extracted | 137,008 |
| # segments after Class2D | N/A |
| # segments after Class3D | 19,995 |
| Resolution (Å) | 3.9 |
| Map sharpening B-factor (Å$^2$) | 0 |
| Helical rise (Å) | 2.41 |
| Helical twist (°) | 179.49 |
| Point Group | $C_1$ |
| **Atomic model** | |
| # non-hydrogen atoms | 2480 |
| # protein residues | 304 |
| R.m.s.d. bonds (Å) | |
| R.m.s.d. angles (°) | |
| Molprobity clashscore, all atoms | 11.1 |
| Molprobity score | 2.1 |
| Poor rotamers (%) | 0 |
| Ramachandran outliers (%) | 0 |
| Ramachandran allowed (%) | 9.0 |
| Ramachandran favored (%) | 91.0 |
| Cβ deviations > 0.25 Å (%) | 0 |
| Bad bonds (%) | 0 |
| Bad angles (%) | 0 |
